# Connectome Gradient Dysfunction in Major Depression and Its Association with Gene Expression Profiles

**DOI:** 10.1101/2020.10.24.352153

**Authors:** Mingrui Xia, Jin Liu, Andrea Mechelli, Xiaoyi Sun, Qing Ma, Xiaoqin Wang, Dongtao Wei, Yuan Chen, Bangshan Liu, Chu-Chung Huang, Yanting Zheng, Yankun Wu, Taolin Chen, Yuqi Cheng, Xiufeng Xu, Qiyong Gong, Tianmei Si, Shijun Qiu, Ching-Po Lin, Jingliang Cheng, Yanqing Tang, Fei Wang, Jiang Qiu, Peng Xie, Lingjiang Li, Yong He, DIDA-Major Depressive Disorder Working Group

## Abstract

**Background:** Patients with major depressive disorder (MDD) exhibit concurrent deficits in sensory processing and high-order cognitive functions such as self-awareness and rumination. Connectome mapping studies have suggested a principal primary-to-transmodal gradient in functional brain networks, supporting the spectrum from sensation to cognition. However, whether this principal connectome gradient is disrupted in patients with MDD and how this disruption is associated with gene expression profiles remain unclear.

**Methods:** Using a large cohort of resting-state functional magnetic resonance imaging data from 2,234 participants (1,150 patients with MDD and 1,084 healthy controls) recruited at 10 sites, we investigated MDD-related alterations in the principal connectome gradient. We further used Neurosynth and postmortem gene expression data to assess the cognitive functions and transcriptional profiles related to the gradient alterations in MDD, respectively.

**Results:** Relative to controls, patients with MDD exhibited abnormal global topography of the principal primary-to-transmodal gradient, as indicated by reduced explanation ratio, gradient range, and gradient variation (Cohen’s *d* = −0.16∼-0.21). Focal alterations of gradient scores were mostly in the primary systems involved in sensory processing and in the transmodal systems implicated in high-order cognition. The transcriptional profiles explained 53.9% of the spatial variance in the altered gradient patterns, with the most correlated genes enriched in transsynaptic signaling and calcium ion binding.

**Conclusions:** These results highlight the dysfunction of the core connectome hierarchy in MDD and its linkage with gene expression profiles, providing insights into the neurobiological and molecular genetic underpinnings of sensory-cognitive deficits in this disorder.

## Introduction

Major depressive disorder (MDD) is one of the most common and burdensome psychiatric disorders globally (1). In addition to clinical symptoms including low mood, loss of interest and fatigue, neuropsychological studies suggest that patients with MDD present with deficits in low-level sensory processing and high-order cognitive functions such as self-awareness, rumination, and reward processing (2-5). Although many prior studies have reported widespread abnormalities in brain structure and function in MDD (6-11), the neurobiological mechanism underlying these deficits in low-level sensory processing and high-order cognitive functions remains to be elucidated.

Hierarchical architecture is one of the fundamental organizational principles of the human brain, allowing for information encoding and integration from sensation to cognition (12). Resting-state functional magnetic resonance imaging (R-fMRI) (13) and the gradient decomposition framework (14) enable researchers to noninvasively investigate the hierarchical architecture of the macroscale functional connectome *in vivo* (14, 15). In healthy adults, the network architecture of the macroscale connectome follows a principal gradient along the axis from primary to transmodal systems. Such a pattern provides insights into the neural basis of the spectrum from sensory to cognitive processing (14) and is largely comparable to cortical microstructural myelination (16). Moreover, the principal primary-to-transmodal gradient changes across brain development (17) and is altered in brain disorders such as autism (18). In patients with MDD, R-fMRI studies have revealed alterations in functional activity and connectivity involving the primary visual and sensorimotor systems (6, 19-21) and transmodal systems such as the default mode network (DMN) and frontoparietal network (FPN) (19, 20, 22-24). However, no studies have reported whether and how the principal primary-to-transmodal gradient of the functional connectome is disrupted in this clinical population. The characterization of the principal connectome gradient in MDD would provide insights into the hierarchical network mechanisms underlying the interplay between sensory and high-order cognitive processing in MDD patients.

Notably, much research has indicated that MDD is a moderately heritable disorder (25). Genome-wide association studies (GWAS) have identified several risk variants of genes linked to MDD, and some of the robustly identified genes play key roles in the biological functions of presynaptic differentiation and neuroinflammation (26). To date, measuring gene expression in brain tissue *in vivo* has been extremely difficult. The integration of gene expression profiles in the postmortem brain with connectomes derived from neuroimaging data provides an unprecedented opportunity to bridge the gap between the microlevel transcriptome profile and the macroscale brain network (27-30). The functional architectures of brain connectomes (e.g., network hubs) are associated with gene expression profiles involving ion channel activity and oxidative metabolism (28, 31). Distinct gene expression profiles can also explain the variances in the spatial patterns of alterations in brain structures in different psychiatric states, including schizotypy (32) and autism (33). Therefore, if patients with MDD exhibit disturbances in the macroscale connectome gradient, we speculate that these functional brain abnormalities might be associated with the transcriptome profiles. The elucidation of such an association would enhance our understanding of the molecular genetic underpinnings of the dysfunctional connectome hierarchy in MDD.

To address these gaps in understanding, in the present study we employed a large multisite R-fMRI dataset from 2,234 individuals and postmortem gene expression data from the brains of six donors from the Allen Institute for Brain Science (AIBS) (27). We investigated the functional connectome gradients in MDD and established their associations with the transcriptome profile. Specifically, we hypothesized that (*i*) the principal primary-to-transmodal gradient is disrupted in patients with MDD, where abnormalities exist in both the global gradient topography and the focal gradient scores of primary and transmodal systems; (*ii*) the regions with an altered connectome gradient in MDD are associated with multiple functional domains, including low-level sensory processing, such as somatosensory and visual perception, and high-order cognitive functions, such as self-referential processing and theory of mind; and (*iii*) the spatial patterns of MDD-related gradient alterations are associated with gene expression profiles that are enriched in particular biological processes (e.g., synapse-related functions).

## Materials and Methods

### Imaging Dataset and Preprocessing

This study included 2,414 participants (1,276 patients with MDD and 1,138 controls) who were recruited from 10 research centers through the Disease Imaging Data Archiving -Major Depressive Disorder Working Group (DIDA-MDD). All patients were diagnosed according to the Diagnostic and Statistical Manual of Mental Disorders IV (DSM-IV) criteria for MDD (34) and did not meet the criteria for any other Axis I disorders. The severity of depression was rated using the Hamilton depression rating scale (HDRS) (35). The controls did not have a current or lifetime history of any Axis I disorder. The exclusion criteria for all participants included MRI contraindications, a history of drug or alcohol abuse, concomitant major medical disorders, head trauma with consciousness disturbances, or any neurological disorders. After strict quality control for both clinical and imaging data (see Supplement), the final sample included 2,234 participants (1,150 patients with MDD and 1,084 controls, Table 1). The study was approved by the ethics committees of each research center, and written informed consent was obtained from each participant.

**Table 1.** Demographic and clinical characteristics of participants.

| Center | Group | Age, mean (SD), yr | Sex (M/F) | Duration of illness, mean (SD), yr | Medicated (Yes/No) | HDRS*, mean (SD) | Mean FD, mean (SD), mm |
| --- | --- | --- | --- | --- | --- | --- | --- |
| CMU, Shenyang | Patient (N=125) | 27.91 (9.70) | 39/86 | 1.65 (3.17) | 49/76 | 21.44 (8.67) | 0.115 (0.072) |
|  | Controls (N=249) | 27.24 (8.20) | 103/146 |  |  |  | 0.107 (0.057) |
| | *t* or $\chi^2/P$ | 0.70/0.484 | 3.33/0.068 | | | | 1.07/0.286 |
| CSU, Changsha | Patient (N=177) | 36.28 (10.21) | 77/100 | 2.52 (3.83) | N.A. | 31.39 (7.82) | 0.141 (0.073) |
|  | Controls (N=108) | 32.31 (7.96) | 62/46 |  |  |  | 0.134 (0.064) |
| | *t* or $\chi^2/P$ | 3.45/0.001 | 5.19/0.023 | | | | 0.88/0.382 |
| GCMU1, Guangzhou | Patient (N=34) | 29.41 (8.27) | 9/25 | 0.65 (0.70) | 0/34 | 21.85 (2.25) | 0.094 (0.030) |
|  | Controls (N=34) | 30.09 (10.88) | 10/24 |  |  |  | 0.096 (0.033) |
| | *t* or $\chi^2/P$ | -0.29/0.774 | 0.07/0.787 | | | | -0.32/0.750 |
| GCMU2, Guangzhou | Patient (N=66) | 29.48 (9.91) | 25/41 | 0.76 (1.00) | 0/66 | 22.30 (3.57) | 0.089 (0.057) |
|  | Controls (N=66) | 29.33 (10.12) | 31/35 |  |  |  | 0.086 (0.042) |
| | *t* or $\chi^2/P$ | 0.29/0.774 | 1.12/0.291 | | | | 0.29/0.770 |
| KMU, Kunming | Patient (N=41) | 34.20 (9.37) | 20/21 | 1.13 (1.28) | N.A. | 23.61 (4.64) | 0.186 (0.083) |
|  | Controls (N=50) | 39.72 (11.97) | 28/22 |  |  |  | 0.167 (0.064) |
| | *t* or $\chi^2/P$ | -2.47/0.015 | 0.47/0.492 | | | | 1.26/0.211 |
| PKU, Beijing | Patient (N=75) | 31.51 (7.86) | 44/31 | 0.52 (0.47) | 0/75 | 25.35 (4.77) | 0.175 (0.063) |
|  | Controls (N=73) | 31.90 (9.01) | 42/31 |  |  |  | 0.185 (0.067) |
| | *t* or $\chi^2/P$ | -0.29/0.775 | 0.02/0.889 | | | | -0.91/0.363 |
| SCU, Chengdu | Patient (N=50) | 34.44 (12.90) | 25/25 | 1.17 (1.60) | 25/25 | 22.88 (4.25) | 0.111 (0.067) |
|  | Controls (N=41) | 34.83 (17.69) | 17/24 |  |  |  | 0.122 (0.072) |
| | *t* or $\chi^2/P$ | -0.12/0.904 | 0.66/0.416 | | | | -0.71/0.479 |
| SWU, Chongqing | Patient (N=282) | 38.74 (13.65) | 99/183 | 4.20 (5.52) | 124/125 | 20.78 (5.88) | 0.125 (0.054) |
|  | Controls (N=254) | 39.65 (15.80) | 88/166 |  |  |  | 0.134 (0.063) |
| | *t* or $\chi^2/P$ | -0.72/0.472 | 0.01/0.911 | | | | -1.68/0.094 |
| YMU, Taipei | Patient (N=105) | 57.05 (16.21) | 63/42 | 1.21 (1.54) | 79/26 | 11.66 (6.99) | 0.139 (0.082) |
|  | Controls (N=109) | 51.12 (11.70) | 69/40 |  |  |  | 0.128 (0.058) |
| | *t* or $\chi^2/P$ | 3.06/0.003 | 0.25/0.619 | | | | 1.17/0.243 |
| ZZU, Zhengzhou | Patient (N=195) | 18.40 (5.54) | 97/98 | 1.28 (1.48) | 0/195 | 22.43 (5.71) | 0.100 (0.045) |
|  | Controls (N=100) | 22.43 (4.49) | 47/53 |  |  |  | 0.088 (0.039) |
| | *t* or $\chi^2/P$ | -6.29/<0.001 | 0.20/0.655 | | | | 2.16/0.032 |
| All data | Patient (N=1150) | 33.78 (14.99) | 477/673 | 2.10 (3.60) | 277/622 |  | 0.125 (0.067) |
|  | Controls (N=1084) | 34.00 (13.87) | 468/616 |  |  |  | 0.123 (0.063) |
| | *t* or $\chi^2/P$ | -0.37/0.712 | 0.66/0.418 | | | | 0.72/0.475 |

Abbreviations: SD, standard deviation; M, male; F, female; HDRS, Hamilton depression rating scale; FD, framewise displacement; CMU, China Medical University; CSU, Central South University; GCMU, Guangzhou University of Chinese Medicine; KMU, Kunming Medical University; PKU, Peking University; SCU, Sichuan University; SWU, Southwest University; YMU, National Yang-Ming University; ZZU, Zhengzhou University; N.A., not available.
\*The 17-item HDRS was used in the research centers of CMU, GCMU, KMU, PKU, SCU, SWU and ZZU. The 21-item HDRS was used in the research center of YMU. The 24-item HDRS was used in the research center of CSU.

The R-fMRI data were obtained from all participants using 3.0-T MRI scanners with gradient-echo planar imaging sequences. During scanning, the participants were instructed to keep their eyes closed without falling asleep and to move as little as possible. Detailed scanning parameters for each research center are listed in Table S1. R-fMRI data preprocessing was conducted using a standard pipeline as described in our previous work (6). See Supplement for details.

### Gene Expression Dataset and Preprocessing

The microarray-based gene expression data were downloaded from the AIBS website (27). The tissue samples in this dataset were collected from the brains of six adult donors (mean age: 42.5 years, 1 female). Each postmortem hemisphere of the brain was dissected into approximately 500 anatomically discrete samples. Each sample was spatially registered to the Montreal Neurological Institute (MNI) coordinate space according to the T1-weighted images obtained before dissection. Normalization processes were conducted by the AIBS to minimize the potential effects of nonbiological biases and ensure that the gene expression data were comparable among samples within and across the brains. We performed preprocessing for the gene expression microarray data according to a recommended pipeline, including mapping samples to a cortical parcellation with 360 regions (36), probe reannotation and selection, and normalization across donors (37). See Supplement for details.

### Connectome Gradient Analysis

For each individual, we first constructed the voxelwise functional network (18,933 nodes) and then applied the diffusion map embedding approach (14, 18) to estimate the connectome gradient. Briefly, the top 10% of the connections were retained for each node, and cosine similarity was calculated between each pair of nodes. The similarity matrix was further scaled into a normalized angle matrix to avoid negative values (17, 38). Then, diffusion map embedding was applied to capture the gradient components that explain the variance in the connectivity pattern of the functional connectome. The resultant gradient maps were further aligned across individuals using iterative Procrustes rotation (18). The joint embedding alignment (39) was also applied for validation purposes. For each gradient map, we calculated three global topographic metrics including the explanation ratio (i.e., the percentage of connectome variance accounted for by a given gradient), the gradient range (i.e., the difference between the greatest positive and negative regional gradient scores), and the gradient variation (i.e., the standard deviation of the gradient scores). Finally, we utilized ComBat harmonization to correct for site effects on the gradient maps and metrics (6, 40). The between-group differences in the connectome gradient were assessed by using a general linear model (dependent variable, gradient metric; independent variable, group) with age and sex as covariates. For global gradient metrics, the statistical significance threshold was set to *P* < 0.05. For regional gradient score maps, the significance threshold was set to *P* < 0.001 at the voxel level, followed by Gaussian random field (GRF) correction at the cluster level of *P* < 0.05 (41).

### Association Analysis Between Cognitive Functions and Gradient Alterations in MDD

We used Neurosynth (https://neurosynth.org/) (42) to assess the cognitive functions associated with the alterations in connectome gradients in MDD. For each gradient component, the thresholded *Z*-maps derived from the between-group comparisons of regional gradient scores were first divided into MDD-positive (i.e., MDD > controls) and MDD-negative (i.e., MDD < controls) maps. The resultant maps were then analyzed using the “decoder” function of the Neurosynth website. The cognitive terms were visualized on a word-cloud plot with the font size scaled according to their correlation with corresponding meta-analytic maps generated by Neurosynth.

### Association Analysis Between Gene Expression and Gradient Alterations in MDD

We used partial least squares (PLS) regression (43) to explore the association between transcriptional profiles and alterations in regional connectome gradients in MDD. PLS regression can define several components, each of which is a linear combination of the predictor variables that can explain most of the variance in the response variables. Here, we first aligned the gene expression data (10,027 genes) and between-group difference *Z*-map of the principal gradient to a cortical parcellation atlas (36). In our PLS model, the gene expression data were set as the predictor variables, and the *Z*-maps of between-group differences of the principal gradient were set as the response variables. To determine whether the PLS components can significantly explain the variation in the response variables, we adopted a permutation test in which the spatial autocorrelations were corrected by generative modeling (44) to examine whether the *R*^2^ of the PLS component was significantly greater than that expected by chance. Then, for each significant component, we used a bootstrapping method to assess the estimation error of the weight of each gene and further divided the weight by the estimated error to obtain the corrected weight of each gene (45). We ranked the genes according to their corrected weights, which represent their contribution to the PLS regression component. That is, genes were ranked in descending order, beginning with those with the most positive relationship and ending with those with the most negative relationship. Both the descending and ascending sequences were enrolled for the following gene enrichment analysis. The Gene Ontology enrichment analysis and visualization tool (GOrilla, http://cbl-gorilla.cs.technion.ac.il/) (46) was used to identify the enriched Gene Ontology terms of the ranked genes. Consistent with previous studies (33, 45), we used the Benjamini-Hochberg false discovery rate (FDR)-corrected *q*-value < 0.05 to determine statistically significant enrichment. See Supplement for details.

### Effects of Clinical Factors

To investigate the effects of categorized clinical factors on the connectome gradient, we classified the patients into different pairs of subgroups according to their clinical information, including patients with an onset age lower or equal to 21 years vs. higher than 21 years (8), patients suffering from their first episode vs. recurrent episodes, and patients receiving medication vs. not receiving medication. Then, we compared the gradient metrics between each corresponding pair of subgroups by using general linear models.

### Validation Analysis

We validated our results by considering several potential confounding factors. First, we used a leave-one-site-out cross-validation strategy to examine whether our findings were influenced by specific sites. This was implemented by repeating the between-group comparisons on the data, excluding one site at a time. Second, some of the participants were younger than 18 years, which might explain the between-group differences in brain development. Thus, we repeated the statistical analysis for only adult participants (1,002 patients with MDD and 1,034 HCs). Third, to further control for the effect of head motion on R-fMRI connectivity measures, we repeated the between-group comparisons with the mean framewise displacement as an additional covariate. Finally, the use of a joint embedding framework may increase the alignment of individual connectome gradient maps compared to the Procrustes rotation (39). Thus, we reperformed the alignment using joint embedding and then repeated the statistical analysis.

### Data Availability

The core analysis code and result data are publicly available at github.com/mingruixia/MDD_ConnectomeGradient.

## Results

The principal primary-to-transmodal gradient explained 11.9% ± 3.1% of the total connectivity variance (MDD, 11.7% ± 3.1%; HC, 12.1% ± 3.0%, Figure S1), which was organized along a gradual axis from the primary visual/sensorimotor networks (VIS/SMN) to the DMN (Figure 1A), replicating the recent observation of connectome gradients from the primary to the transmodal cortices in healthy adults (14). The spatial patterns of the group-averaged principal gradient maps were remarkably similar between the MDD and HC groups (Spearman’s *ρ* = 0.999, *P* < 0.0001, permutation tests with spatial autocorrelation corrected) (Figure S2). Visual inspection of the histogram revealed that the extremes of the primary-to-transmodal gradient were contracted in MDD relative to the control range (Figure 1B and C). Here, we mainly report the results from the principal connectome gradient. The results from the second and third gradients can be found in Figures S3, S4, and S5 and Tables S2, S3, and S4.

**Figure 1.**
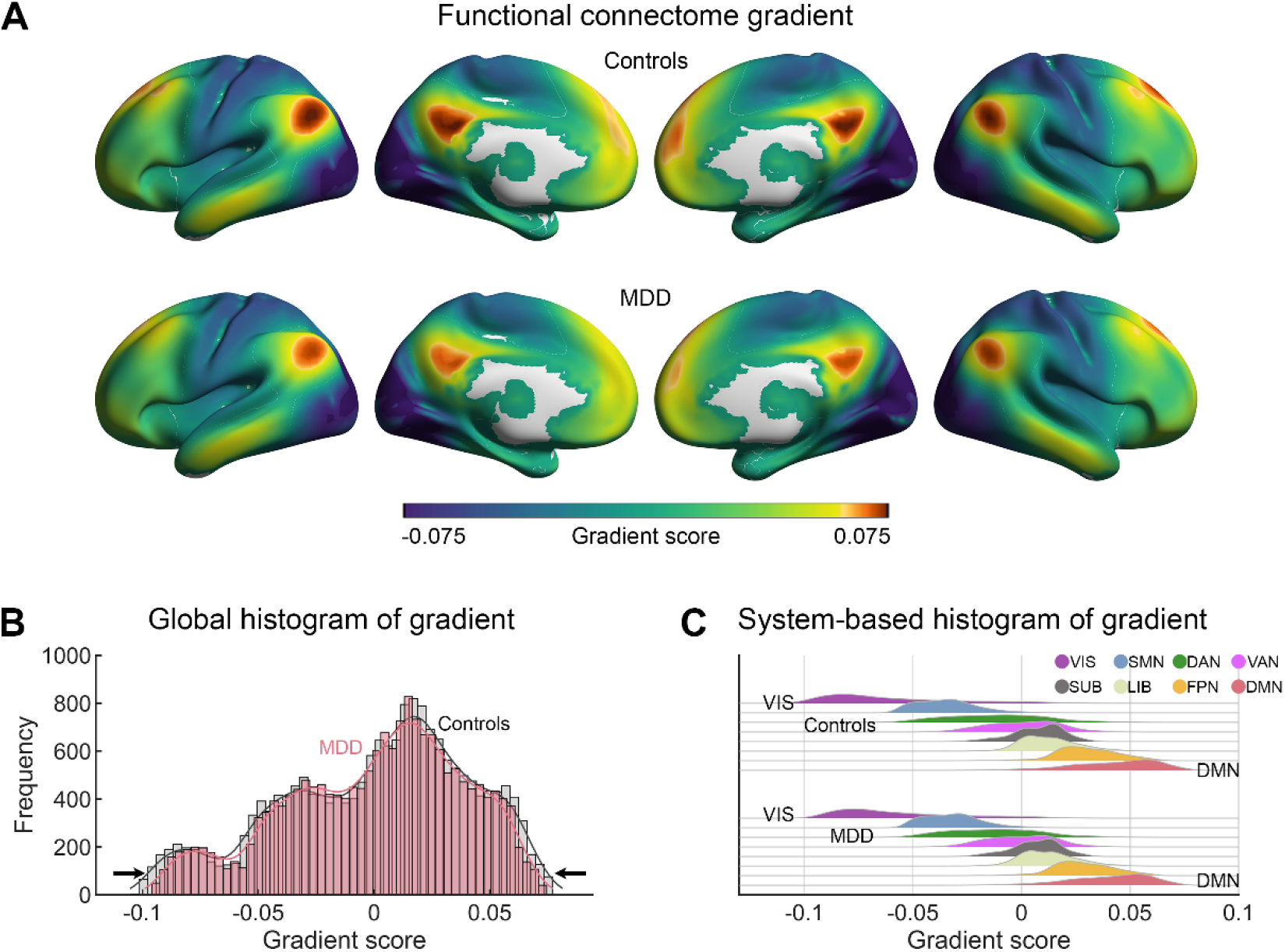
Connectome gradient mapping in patients with MDD and controls. **(A)** The principal gradient was organized along a gradual axis from the primary visual/sensorimotor networks to the default mode network. **(B)** Global and **(C)** system-based histograms show that the extreme values were contracted in patients with MDD relative to healthy controls. Surface rendering was generated using BrainNet Viewer (www.nitrc.org/projects/bnv/)(75) with the inflated cortical 32K surface (36). VIS, visual network; SMN, sensorimotor network; DAN, dorsal attention network; VAN, ventral attention network; SUB, subcortical regions; LIB, limbic network; FPN, frontoparietal network; DMN, default mode network.

### Alterations of Connectome Gradients in MDD

Between-group statistical comparisons showed that the primary-to-transmodal gradient explained less variance in the functional connectome in the MDD group than in the HC group (Cohen’s *d* = −0.16, *P* = 0.0002, Figure 2A and Table S2), suggesting a downgraded status of the hierarchal organization in MDD. Moreover, the patients with MDD showed a narrower range of scores (*d* = −0.21, *P* = 0.000001) and less spatial variation (*d* = −0.20, *P* = 0.000003) than the HCs (Figure 2A and Table S2), indicating a contracted connectome hierarchy in MDD. Regionally, the MDD group showed lower gradient scores in the DMN but higher scores in the VIS and SMN than the HC group (voxel-level *P* < 0.001, GRF-corrected *P* < 0.05) (Figure 2B and Table 2). The identified clusters, particularly the regions in the DMN, VIS, and SMN, showed a strong shift from the periphery to the center in gradient space, indicating the less-specialized connectivity profiles among these regions in MDD (Figure 2C).

**Table 2.** Clusters with significant between-group differences in the principal primary-to-transmodal gradient

| No. | Region | x | y | z | t | Cohen's *d* | Size (mm<sup>3</sup>) |
| --- | --- | --- | --- | --- | --- | --- | --- |
| MDD > Controls |  |  |  |  |  |  |  |
| 1 | Right fusiform/lingual gyri, BA37 | 24 | -48 | -12 | 5.69 | 0.24 | 35968 |
| 2 | Bilateral postcentral gyri, BA3 | -20 | -40 | 68 | 4.53 | 0.19 | 12608 |
| 3 | Left fusiform/lingual gyri, BA19 | -24 | -60 | -16 | 4.91 | 0.21 | 11840 |
| 4 | Left Rolandic operculum, BA13 | -40 | -28 | 20 | 5.60 | 0.24 | 9856 |
| 5 | Right Rolandic operculum, BA13 | 64 | -4 | 12 | 5.20 | 0.22 | 7488 |
| 6 | Right precentral gyrus, BA4 | 44 | -16 | 40 | 4.88 | 0.21 | 3392 |
| 7 | Right middle occipital gyrus, BA19 | 44 | -80 | 8 | 4.30 | 0.18 | 1728 |
| MDD < Controls |  |  |  |  |  |  |  |
| 8 | Bilateral middle frontal gyri/superior frontal gyri, medial part, BA6 | -32 | 8 | 52 | -5.83 | -0.25 | 51136 |
| 9 | Bilateral posterior cingulate gyri/precuneus, BA31 | -8 | -44 | 28 | -5.54 | -0.23 | 13824 |
| 10 | Left angular gyrus, BA39 | -44 | -60 | 28 | -5.29 | -0.22 | 8000 |
| 11 | Right supramarginal gyrus, BA40 | 48 | -40 | 40 | -4.54 | -0.19 | 2816 |
| 12 | Right thalamus | 4 | -12 | 0 | -4.20 | -0.18 | 2304 |

**Figure 2.**
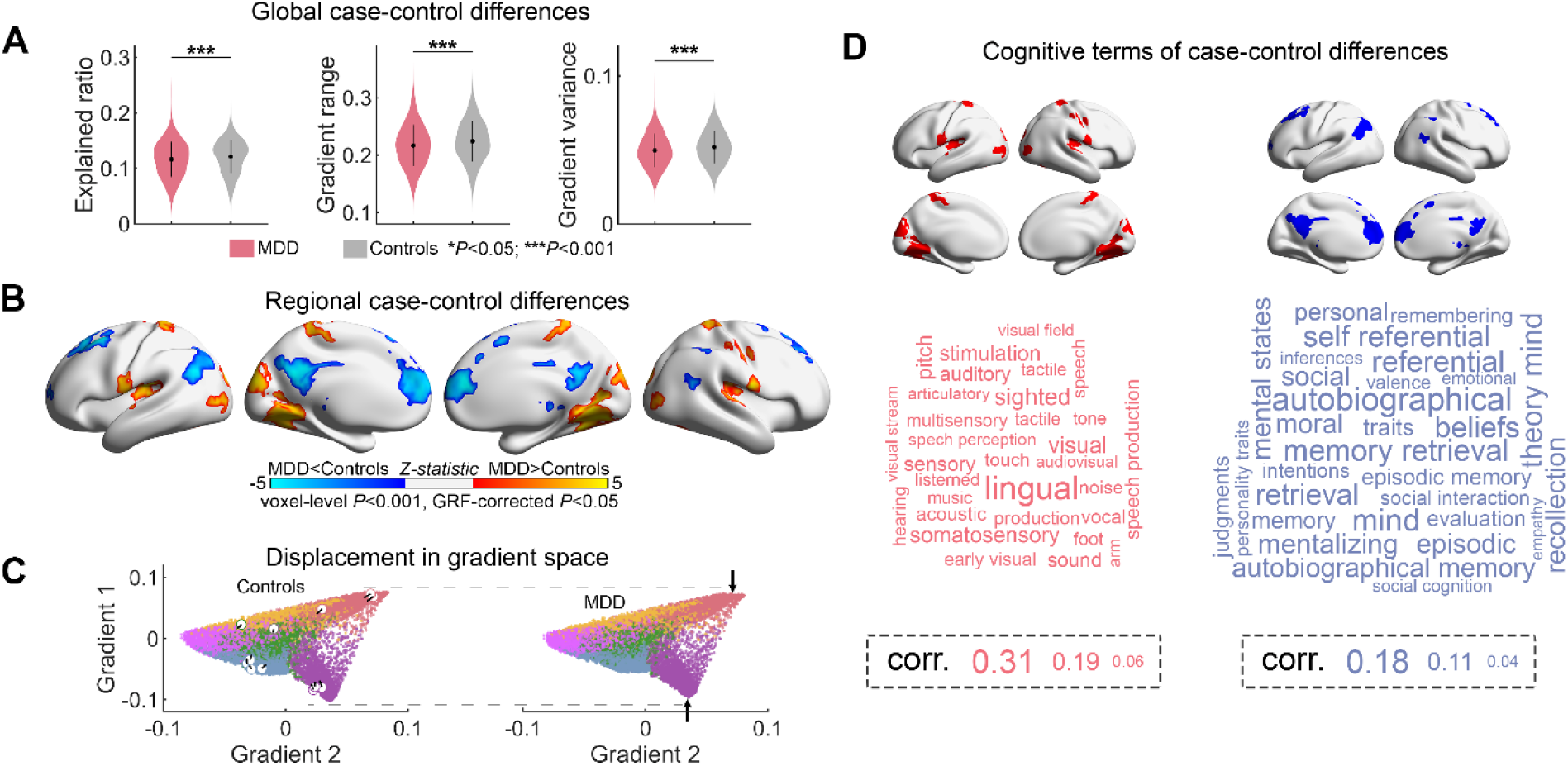
Statistical comparison of gradient metrics. **(A)** Case-control differences in global gradient metrics. ***, *P* < 0.001. **(B)** Voxelwise statistical comparisons between healthy controls and patients with MDD, with higher/lower values in MDD presented as warm/cold colors. The statistical significance level was set as voxel-level *P* < 0.001 and Gaussian random field cluster level-corrected *P* < 0.05. **(C)** Scatter plot for the first two gradients in healthy controls and patients with MDD illustrates a contracted distribution of gradient scores in MDD. Each dot represents a voxel, and its color indicates the corresponding system. The circles represent the peak of the clusters with case-control differences, and their arrows indicate their displacement in gradient space. **(D)** Word clouds of cognitive functions associated with brain regions that exhibited higher (red) or lower (blue) gradient scores in MDD. The font size of the cognitive terms corresponds to the strength of the correlation of corresponding meta-analytic maps generated by Neurosynth.

### Cognitive Functions Relating to Gradient Alterations in MDD

We used Neurosynth to decode the between-group differences in the gradient scores of the primary-to-transmodal component against cognitive functions. We found that the regions with higher gradient scores in MDD were mainly involved in sensorial and perceptional processes, such as visual, somatosensory, and acoustic processes, and regions with lower gradient scores were implicated in DMN-related functions including self-referential, theory of mind, memory retrieval, and autobiographical memory (Figure 2D). These results indicate that the alterations in the principal primary-to-transmodal gradient in MDD are related primarily to both sensory processing and high-level cognitive functions.

### Gene Expression Profiles Relating to Gradient Alterations in MDD

PLS regression was applied to investigate the relationship between the between-group *Z*-map of the principal primary-to-transmodal gradient and gene expression profiles. The first two components of the PLS regression explained 53.9% of the variance in the MDD-related alterations in the principal gradient (*P* < 0.0001 for component 1 and *P* = 0.004 for component 2, correcting for spatial autocorrelations by permutation test, Figure S6). Component 1 represented a transcriptional profile characterized by high expression mainly in the posterior parietal-occipital areas but low expression in prefrontal areas (Figure 3A). Component 2 revealed a gene expression profile with high expression mainly in the sensorimotor, visual and temporal cortices but low expression in the frontoparietal cortices (Figure 3A). The regional mapping of these two components positively correlated with the *Z*-map of the primary-to-transmodal gradient map (component 1: *r* = 0.576, *P* < 0.0001; component 2: *r* = 0.456, *P* < 0.0001, correcting for spatial autocorrelations by permutation test, Figure 3B). The Gene Ontology enrichment analysis revealed that the genes ranked in ascending order of the PLS component 1 weight were enriched in biological processes related to transsynaptic signaling and molecular function of calcium ion binding (FDR-corrected *q* < 0.05, Figure 3C and Table S5). The genes ranked according to the weight of component 2 did not show statistically significant enrichment.

**Figure 3.**
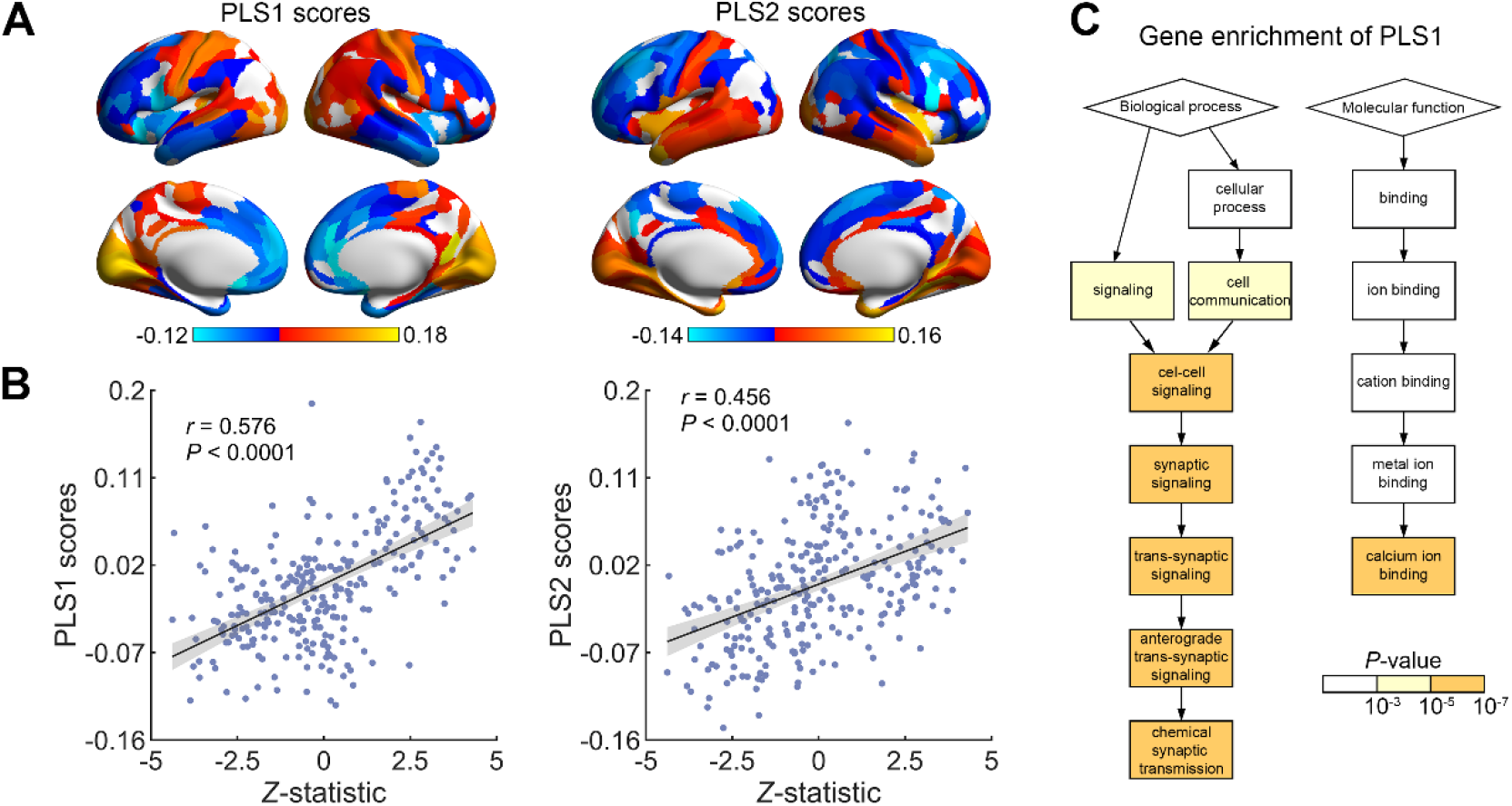
Association between MDD-related gradient alterations and gene expression. **(A)** The first PLS component (PLS 1) identified a gene expression profile with high expression mainly in the posterior parietal-occipital areas but low expression in prefrontal areas. The second PLS component (PLS 2) revealed a gene expression profile with high expression mainly in the sensorimotor, visual and temporal cortices but low expression in the frontoparietal cortices. **(B)** The transcriptional profiles were positively correlated with the between-group *Z*-map of the principal gradient (permutation tests with spatial autocorrelation corrected, 10,000 times). The shadow indicates 95% confidence intervals. Each dot represents a region. **(C)** Genes ranked in ascending order of PLS 1 weight were enriched in the biological process of transsynaptic signaling and molecular function of calcium ion binding (FDR _*q*_ < 0.05).

### Clinical Relations to Connectome Gradients in MDD

Patients with an onset in adolescence (age ≤ 21 years, N = 303) showed a narrower gradient range (*d* = −0.28, *P* = 0.001) and a smaller region variation (*d* = −0.18, *P* = 0.026) of the principal primary-to-transmodal gradient than patients who had an onset age older than 21 years (N = 293) (Figure 4). There were no statistically significant differences in the topographic features of the principal gradient between patients who were and were not taking medication or patients in their first episode and recurrent patients (Table S6). Voxelwise comparisons showed that there was no statistically significant difference in regional gradient scores between any of these clinical category pairs after correcting for multiple comparisons.

**Figure 4.**
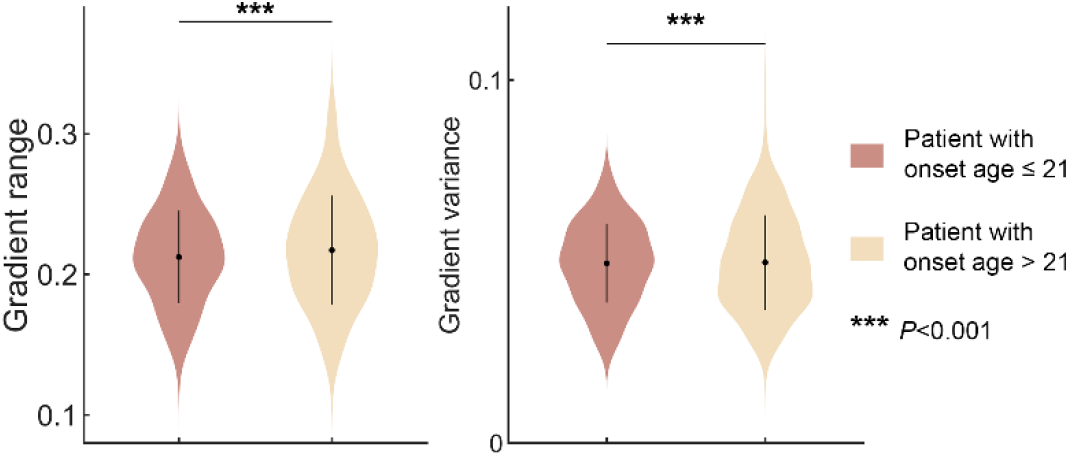
Effects of onset age on gradient topography. The range and variance of the principal gradient were significantly lower in patients with an onset age ≤ 21 years than in those with an onset age > 21 years.

### Validation Results

Overall, the MDD-related alterations in the principal connectome gradient in different validation strategies remained highly similar to our main findings (Figure S7 and S8, and Table S7 and S8).

## Discussion

Using a large cohort of R-fMRI data, we demonstrated connectome gradient dysfunction in patients with MDD, including disrupted global topography and focal alterations of gradient scores in the primary areas involved in sensory processing and transmodal areas involved in high-order cognitive processing. These gradient changes are tightly associated with transcriptional profiles, and the most correlated genes are enriched in transsynaptic signaling and calcium ion binding. These findings provide insights into the understanding of the neurobiological mechanisms underlying sensory-cognitive deficits in patients with MDD.

### Downgraded and Contracted Connectome Hierarchy in MDD

We demonstrated less-explained variance in the functional connectome and a narrower distribution range of the principal primary-to-transmodal gradient in MDD, which suggests a downgraded and contracted connectome hierarchy. Disturbances in the functional architecture of the macroscale brain network have recently been considered critical in the pathology of depression (19, 47, 48). Specifically, network analysis based on graph theory has revealed a replicable pattern towards a randomized configuration of the functional connectomes as characterized by more long-range connections in patients with MDD (21, 49, 50). These hyperconnections break the balance between functional integration and functional segregation and further affect the hierarchical architecture of the brain networks. Notably, such an alteration of the core connectome gradient was not the result of the disconnection of a single system but of widely distributed systems from the low-level primary to the high-order transmodal cortices.

We observed that several specific regions in the SMN/VIS and DMN shifted from the periphery to the center in the hierarchical space, implying a trend towards dedifferentiated connection profiles between these systems. This phenomenon is comparable to previous findings from connectome modular studies in MDD, where hyperconnections were found among the primary systems and the transmodal DMN and FPN (20, 23, 51). In a hierarchical organization, the primary brain systems receive external stimulation signals and process them into abstract representations (52), while the transmodal brain systems integrate the processed information with internal control, memory, and emotion to guide interactions with the external environment (12, 52). Specifically, the primary sensory cortices, including the primary visual, auditory, and somatosensory systems, play a key role in visual, auditory, and tactile perception (53-55).

In contrast, the core regions of the DMN, such as the medial prefrontal cortex, posterior cingulate cortex, and precuneus, play important roles in self-referential, theory of mind, memory retrieval, and autobiographical memory (56-58). The contracted hierarchy involving multilevel systems (i.e., overintegration), therefore, may reflect incomplete or blunt bottom-up information processing from the primary systems and failures in the corresponding top-down processing from the high-order systems, thus resulting in clinical and cognitive impairments in multiple domains in MDD (59). Additionally, the degree of downgrade and contraction in the connectome hierarchy increases as the patient onset age decreases, indicating that an early onset is linked with more severe changes in the connectome hierarchy. Given the close relationship between the pathology of MDD and age, future research focusing on the interactive effect between development and depression on the connectome hierarchy is essentially important.

### Gene Expression Profiles for Transsynaptic Signaling and Calcium Ion Binding in MDD

Our connectome-transcriptome association analysis established a link between MDD-related changes in connectome gradients and gene expression enriched in transsynaptic signaling and calcium ion binding. Transsynaptic signaling is one of the most fundamental biological processes that contributes to a series of critical molecular functions, including instructing the formation of synapses, regulating synaptic plasticity, and matching pre- and postsynaptic neurons (60, 61). It enables the establishment of complex neuronal networks supporting effective information transfer and processing throughout the brain. The variation in synaptic signaling across the cerebral cortex has been demonstrated to be organized along the axis of the cortical hierarchy, which corresponds well to the principal gradient of macroscale functional connectomes (62, 63). Notably, disruptions in transsynaptic signaling in many of the key pathways can influence the formation and stability of synapses and have been known to play roles in the pathology of depression (64). For example, studies in postmortem tissues and rodent models revealed that exposure to chronic stress can disrupt the pathway of brain-derived neurotrophic factor (BDNF)-tropomyosin-related kinase B (TrkB) receptor signaling by reducing the downstream extracellular signal-regulated kinase (ERK) and Akt pathways in the hippocampus and prefrontal cortex (65, 66). Disturbances in these pathways can decrease the expression and function of BDNF and further cause neuronal atrophy in regions that are implicated in depression (67). Consistent with our findings, a recent study combining gene coexpression networks and genome-wide summary statistics also revealed that MDD risk genes were enriched in gene modules involving transsynaptic signaling (68). In addition to transsynaptic signaling, calcium ion binding is another crucial molecular function for intracellular signaling. In particular, calcium ion binding can occur in signal transduction resulting from the activation of ion channels or as a second messenger in wide-ranging physiological pathways involving synaptic plasticity. In MDD, evidence from postmortem studies suggests that the density of calbindin-immunoreactive GABAergic neurons is reduced in the dorsolateral prefrontal cortex of patients (69). Therefore, our findings provide further evidence that the disrupted connectome hierarchy architecture in MDD is associated with the gene expression profile related to these two general molecular mechanisms. However, we were unable to determine whether microlevel transcriptional dysregulation resulted in macrolevel connectome dysfunction or whether either of these were causally influenced by risk factors for MDD, such as environmental risk factors.

### Limitations and Future Research

First, cognitive performance was not measured in patients, which limited the opportunity to study the associations between connectome hierarchy disruptions and different aspects of cognitive domains in MDD. To address this limitation, we examined the associations between the gradient alteration maps and the meta-analytic cognitive function maps from the widely used Neurosynth database (42). Second, longitudinal information such as clinical response to treatment was not included here. Previous studies have suggested that functional brain abnormalities can be normalized after antidepressant (70), electroconvulsive therapy (71), or deep brain stimulation surgery (72) in patients with MDD. Future studies using longitudinal datasets are required to enhance our understanding of the effects of treatment on connectome hierarchy architectures in MDD and to provide imaging biomarkers for evaluating treatment effects. Third, several studies reported an abnormal gradient topography of the cerebral cortex in the autism spectrum (18) and of the cerebellar cortex in schizophrenia (73). Notably, our study showed that patients with MDD exhibited more widespread gradient alterations in the primary sensory cortices. Nonetheless, the specificity of gradient alteration for each disorder remains to be further elucidated. Fourth, the fMRI signal is bound via neurovascular coupling to neuronal processes, thus containing both metabolic and vascular-hemodynamic information. Previous works have suggested metabolic or vascular-hemodynamic alterations in MDD, such as alteration of cerebral blood flow (74). Future studies combining electrophysiological recordings and metabolic data (e.g., PET/ASL) are required to distinguish disruption of neuronal processing from other pathologies in MDD. Finally, the gene expression data from the AIBS were sampled from donors without a diagnosis of MDD. Thus, the observed association between connectome hierarchy and transcriptome profiles should be considered cautiously. Future studies with larger samples of whole-brain genome-wide gene expression data from patients with MDD could provide further evidence. Despite these limitations, our study highlights the dysfunction of the core connectome hierarchy in MDD and its linkage with gene expression profiles, providing insights into the neurobiological and molecular genetic underpinnings of sensory-cognitive deficits in this disorder.

## Acknowledgements and Disclosures

This work was supported by the National Natural Science Foundation of China (81671767, 82071998, 82021004, 81620108016, 91432115, 81630031, 31771231, 31271087, 81271499, 81571311, 81571331, 81571331, 81171286, 91232714, 81621003, 81920108019, 81771344, and 81660237), the Beijing Nova Program (Z191100001119023), Fundamental Research Funds for the Central Universities (2020NTST29), the National Key R&D Program of China (2018YFA0701400), the Changjiang Scholar Professorship Award (T2015027), the National High Tech Development Plan (863) (2015AA020513), the National Science and Technologic Program of China (2015BAI13B02), the National Basic Research Program of China (2013CB835100), the Natural Science Foundation of Chongqing (cstc2019jcyj-msxmX0520), the Medical Science and Technology Research Project of Henan Province (201701011), and Shanghai Science and Technology Innovation Plan (17JC1404105 and 17JC1404101). The authors thank the Allen Institute for Brain Science for providing the gene expression data.

The authors report no biomedical financial interests or potential conflicts of interest.

## Notes

### Competing Interest Statement

The authors have declared no competing interest.

