## Supplement for "Connectome Gradient Dysfunction in Major Depression and Its Association with Gene Expression Profiles"

**SI Appendix for "Connectome Gradient Dysfunction in Major Depression and Its Association with Gene Expression Profiles": Supplemental Methods, Figures, and Tables**

### SI Materials and Methods

#### *Imaging Dataset*

A total of 2,414 participants, including 1,276 patients with major depressive disorder (MDD) and 1,138 healthy controls (HCs), were recruited from 10 research centers in China (China Medical University, CMU; Central South University, CSU; Guangzhou University of Chinese Medicine, GCMU (two datasets); Kunming Medical University, KMU; Peking University Sixth Hospital, PKU; Sichuan University, SCU; Southwest University, SWU; National Yang-Ming University, YMU; and Zhengzhou University, ZZU). All patients were diagnosed according to the Diagnostic and Statistical Manual of Mental Disorders IV (DSM-IV) criteria for MDD (1). The severity of depression was rated using the Hamilton Depression Rating Scale (HDRS) (2). Strict quality control was performed for both clinical and imaging data, and 180 participants were excluded due to a lack of demographic information (N = 5), younger age (7 years old, N = 1), a change in their diagnosis during follow-up interviews (N = 7), duplicate data in the data transfer or errors in the raw DICOM data (N = 10), different scanning parameters or incomplete scans (N = 24), abnormalities in the anatomical brain images (N = 14), excessive head motion (exceeding 3 mm of translational movement, 3° of rotational movement or 0.5 mm of mean framewise displacement, N = 71), incomplete coverage of the entire brain (N = 46), error in normalization (N = 1), and an abnormal temporal signal-to-noise ratio (N = 1). The final sample included 2,234 participants (1,150 patients with MDD and 1,084 HCs, Table S1). The study was approved by the ethics committees of each center, and written informed consent was obtained from each participant. All resting-state functional MRI (R-fMRI) data were obtained on 3.0-T MRI scanners with gradient-echo planar imaging sequences. During the scan, the participants were instructed to keep their eyes closed without falling asleep and to move as little as possible. Detailed scanning parameters for each center are listed in Table S2.

#### *Data Preprocessing*

R-fMRI image preprocessing was conducted with SPM12 (www.fil.ion.ucl.ac.uk/spm/) and SeeCAT (https://github.com/mingruixia/MDD_ConnectomeGradient/tree/main/0.Preprocessing/SeeCAT). Briefly, the first ten time points (the first five time points for the CSU, GCMU1 and ZZU datasets due to their short scan times) were discarded. Subsequent preprocessing steps included slice-timing correction and head-motion correction. Next, the motion-corrected functional images were normalized to the standard space using an echo planar imaging (EPI) template, resampled to 3-mm isotropic voxels, and further smoothed with a 6-mm full-width at half-maximum Gaussian kernel. Linear detrending was performed, and several confounding covariates, including the Friston-24 head-motion parameters and the white matter and cerebrospinal fluid signals, were regressed out from the time series for all voxels. Subsequently, temporal bandpass filtering (0.01-0.08 Hz) was applied. Finally, a "scrubbing" procedure was performed on individual preprocessed datasets to remove outlier data due to head motion (3). Specifically, for volumes with a framewise displacement exceeding a threshold of 0.5 mm, we replaced the volumes and their adjacent volumes (2 forward and 1 backward frames) with linearly interpolated data.

#### *Connectome Gradient Analysis*

We constructed individual functional connectomes at the voxel level. To reduce the computational complexity, we resampled the preprocessed R-fMRI images to a 4-mm isotropic resolution. For each individual, a functional connectivity matrix was first estimated by calculating the Pearson correlation coefficients between each pair of gray matter nodes (18,933 voxels). The top 10% of the connections representing the backbone of the connectome were retained for each node, and the cosine similarity was calculated between each pair of nodes. The similarity matrix was further scaled into a normalized angle matrix to avoid negative values (4, 5). Then, diffusion map embedding was applied to capture the gradient components that could explain the variance in the connectivity pattern of the functional connectome. Following the previous recommendation, we set the manifold learning parameter $\alpha = 0.5$ (4, 6, 7). For each gradient map, the explanation ratios for the connectome variance, gradient range, and gradient variance were calculated. Procrustes rotation was performed to align individual gradient maps across subjects (4, 7). Finally, we utilized the ComBat model, an empirical Bayes-based multivariate linear mixed-effects regression, to correct for site effects on the gradient map and measurements (8, 9). The between-group differences in the gradient measurements were determined by two-sample $t$ test with age and sex controlled. The significance level of the voxelwise comparison was set to a voxel-level $P < 0.001$ with a cluster-level Gaussian random field-corrected $P < 0.05$ (10).

#### *Association Analysis Between Cognitive Functions and Gradient Alterations in MDD*

We used Neurosynth (https://neurosynth.org/) (11) to assess the cognitive functions associated with alterations in the connectome gradients in MDD. The thresholded $Z$-maps derived from the between-group comparisons for each gradient were first divided into MDD-positive (i.e., MDD > controls) and MDD-negative (i.e., MDD < controls) maps. The resultant maps were then uploaded to Neurovault and analyzed using the "decoder" function of the Neurosynth website. For each of the maps, the noncognitive terms (e.g., anatomical and demographic terms) were removed, and the top 30 cognitive terms were selected. The cognitive terms were visualized on a word-cloud plot with the font size scaled according to their correlation with corresponding meta-analytic maps generated by Neurosynth.

#### *Gene Expression Data Preprocessing*

We used whole-brain microarray-based gene expression data from the Human Brain Atlas downloaded from the Allen Institute for Brain Science (AIBS) website (http://human.brain-map.org, RRID: SCR_007416) (12). Human brain tissue samples of this atlas were collected from the brains of six adult donors (mean age: 42.5 years, range: 24-57 years, 1 female), including two complete brains and four left hemispheres. Each postmortem hemisphere of the brain was dissected into approximately 500 anatomically discrete samples. Each sample was spatially registered to the Montreal Neurological Institute (MNI) coordinate space according to the T1-weighted images obtained before dissection, and the locations of all samples are provided in MNI coordinates. Each sample underwent microarray analysis and preprocessing to quantify gene expression across 58,692 probes. Normalization processes were conducted by the AIBS to minimize the potential effects of nonbiological biases and ensure that the gene expression data were comparable among samples within and across the brains. Given that the AIBS dataset did not cover the whole brain at the voxel level, we utilized cortical 360-region brain parcellation (13, 14) to perform gradient-gene expression association analysis. We performed preprocessing for the gene expression microarray data of brain tissue samples by using the Allen Human Brain Atlas (AHBA) processing pipeline (https://github.com/BMHLab/AHBAprocessing) with the default recommended setting (15). This preprocessing procedure is built on a systematic assessment of workflow for combining AHBA and neuroimaging data (15). Briefly, the probe-to-gene annotations were verified using the hg38 sequencing database. Then, probes with values that did not exceed background noise were filtered. The probe with the highest correlation to RNA-seq data was selected to index expression for a gene. Then, each tissue sample was assigned to its nearest cortical region of the 360-parcellation. Samples with a distance greater than 2 mm to any of the 360 regions were excluded. These procedures resulted in 820 brain tissue samples covering 284 regions, with each sample containing the expression of 10,027 genes. Subsequently, a two-step scaled robust sigmoid normalization approach was used to correct for both intersample and intersubject variability. For each sample, normalization was applied across all the probes within the sample. Then, for each subject, normalizations were performed for each probe across all the samples. Finally, for each region, gene expression was obtained by averaging all samples from six donors, resulting in a gene expression map (284 regions × 10,027). Data from four regions were further excluded in the following analysis due to their low overlapping percentage (< 50%) with the gray matter mask used in gradient analysis.

***Association Analysis Between Gene Expression and Gradient Alterations in MDD***

We used partial least squares (PLS) regression to explore the association between transcriptional profiles and alterations in the principal connectome gradient in MDD. Prior to PLS regression, we aligned the gene expression data and between-group difference *Z*-maps to a cortical parcellation atlas with 360 regions (13) due to the sparse distribution of gene expression data. Eighty regions were excluded due to missing data (see Gene Expression Data Preprocessing). PLS regression can define several components, each of which is a linear combination of the predictor variables (i.e., gene expression) that can explain most of the variance in the response variables (i.e., between-group difference *Z*-maps of connectome gradients). The predictor variables matrix $X$ and the response variables matrix $Y$ are first centered, resulting in $X_0$ and $Y_0$, respectively. Component $i$ of the PLS regression is then weighted by $p_i$ and $q_i$ to calculate the component scores $T_i$ and $U_i$ for $X_0$ and $Y_0$, respectively:

$$T_i = X_0 p_i + E;\ \ U_i = Y_0 q_i + F$$

where $E$ and $F$ are error terms. Then, the weight vectors $p_i$ and $q_i$ and the component scores $T_i$ and $U_i$ are estimated to ensure the maximum covariance between $T_i$ and $U_i$. Thus, the regression of the predictor variables and response variables can be defined as follows:

$$U_i \sim T_i \ \text{ or } \ Y_0 q_i = B_{0i} + B_{1i} X_0 p_i + G$$

where $G$ is an error term and $B_{1i}$ and $B_{0i}$ are the regression coefficient and intercept, respectively. The $R^2$ of the fitting for each component illustrates how much the predictive variables can explain the variance in the response variables. Here, in our PLS model, the gene expression data of the brain nodes (280 nodes × 10,027 genes) were set as the predictor variables $X$, and the $Z$ values of the principal gradient (280 nodes × 1 statistics) were set as the response variables $Y$.

A permutation test combined with spatial-autocorrelation correction (16) was performed to determine whether the $R^2$ derived from PLS regression analysis was significantly greater than that expected by chance. In each permutation, the between-group difference $Z$-map of the connectome gradient was first randomly shuffled across voxels. Then, the variogram of the original $Z$-map was estimated and used to smooth and rescale the permuted map, resulting in a spatial autocorrelation-reserving surrogate $Z$-map. Then, the surrogate $Z$ values were used as the response variable in the PLS regression, and the corresponding $R^2$ values were recorded. The permutation was repeated 10,000 times to generate the null model. The real $R^2$ values of the first few components that explained over ten percent of the variance in the response variables (here, only one component over ten percent) were compared with those from this null model to determine whether the real $R^2$ values were significantly greater than those expected by chance. Similarly, the significance levels of spatial correlations between PLS scores and the Z-map were also determined by using permutation tests. Then, for each significant component, we used a bootstrapping method to assess the estimation error of the weight of each gene and further divided the weight by the estimated error to obtain the corrected weight of each gene (17).

We ranked the genes according to their corrected weights, which represent their contribution to the PLS regression components. Both the positive and negative sequences were enrolled for the following gene enrichment analysis. The Gene Ontology enrichment analysis and visualization tool (GOrilla, http://cbl-gorilla.cs.technion.ac.il/) (18) was used to identify the enriched Gene Ontology terms of the ranked genes from each significant component. Specifically, we used a $P$-value threshold of $10^{-5}$ in the advanced parameter settings and applied the Benjamini-Hochberg false discovery rate (FDR) method to correct for the multiple tests. In the main results, Gene Ontology terms with an FDR $q$-value less than 0.05 were reported.

### Supplementary Figures

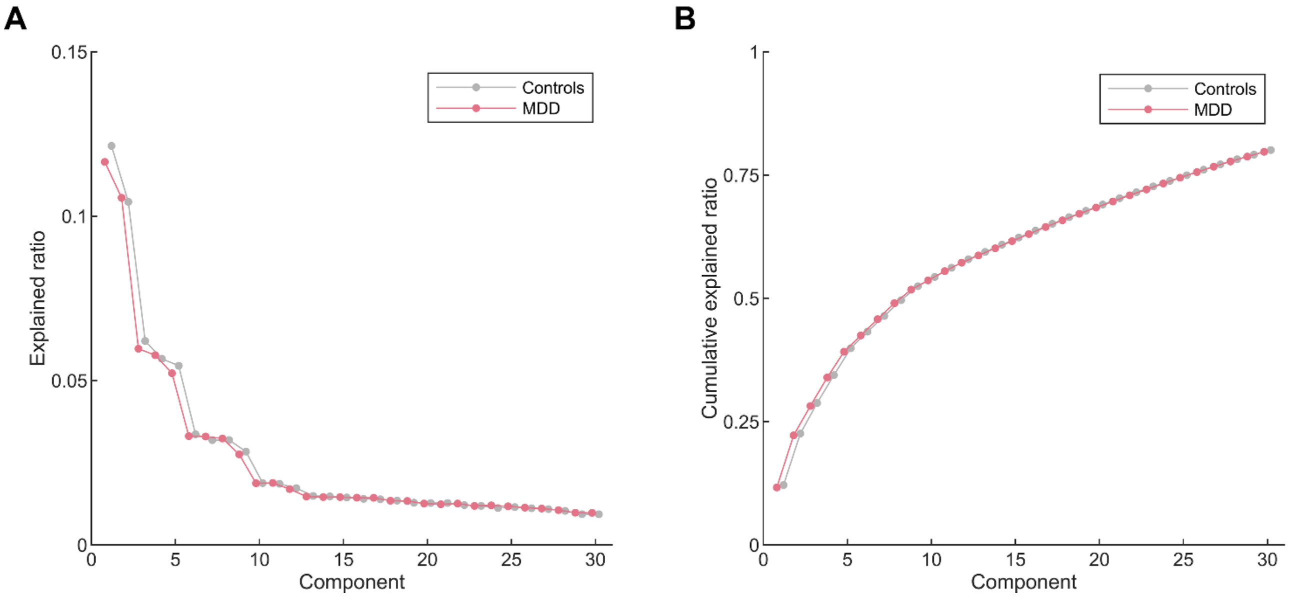


**Figure S1.** (**A**) The averaged explained ratio (λ values) and (**B**) the cumulative averaged explained ratio of the first 30 diffusion embedding components in the controls and MDD groups. Controls are shown in gray and MDD in red. The first three gradients explained 28.5% ± 3.9% of the total variance in the connectome across all individuals (MDD, 28.2% ± 3.9%; HC, 28.9% ± 3.9%). The explained ratios of Gradients 1 and 3 were significantly lower in the MDD group than in the control group (Gradient 1: $P$ = 0.00019; Gradient 3: $P$ = 0.019).

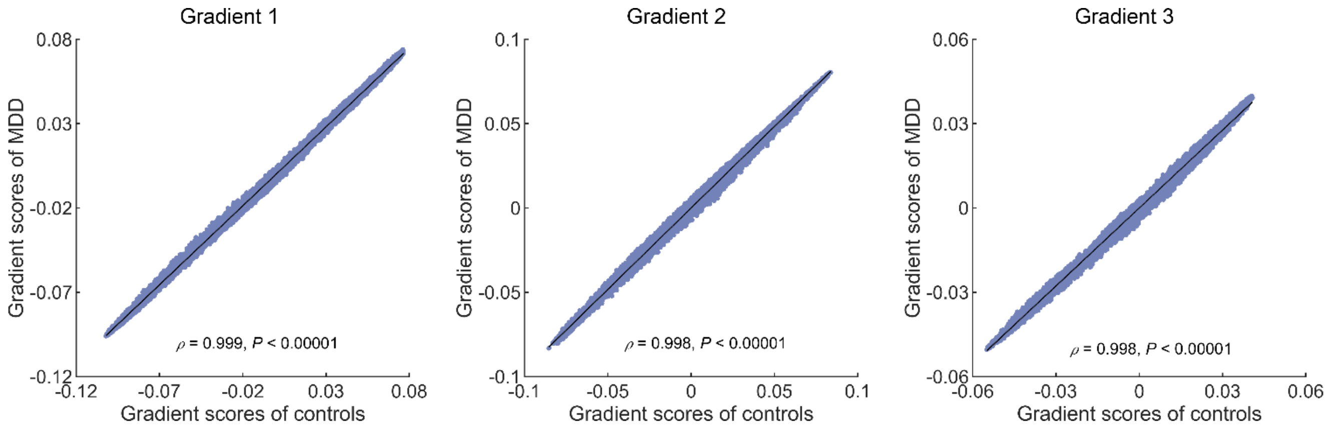


**Figure S2.** Spatial correlations of the group-averaged gradient maps between the controls and MDD groups. Each dot represents a brain voxel. The spatial patterns of the group-averaged gradient maps were remarkably similar between the MDD and control groups, with Spearman's $\rho = 0.999$, 0.998, and 0.998 for Gradients 1, 2, and 3, respectively (all $P < 0.0001$). These correlations were corrected for spatial autocorrelations by using a permutation test (N = 10,000).

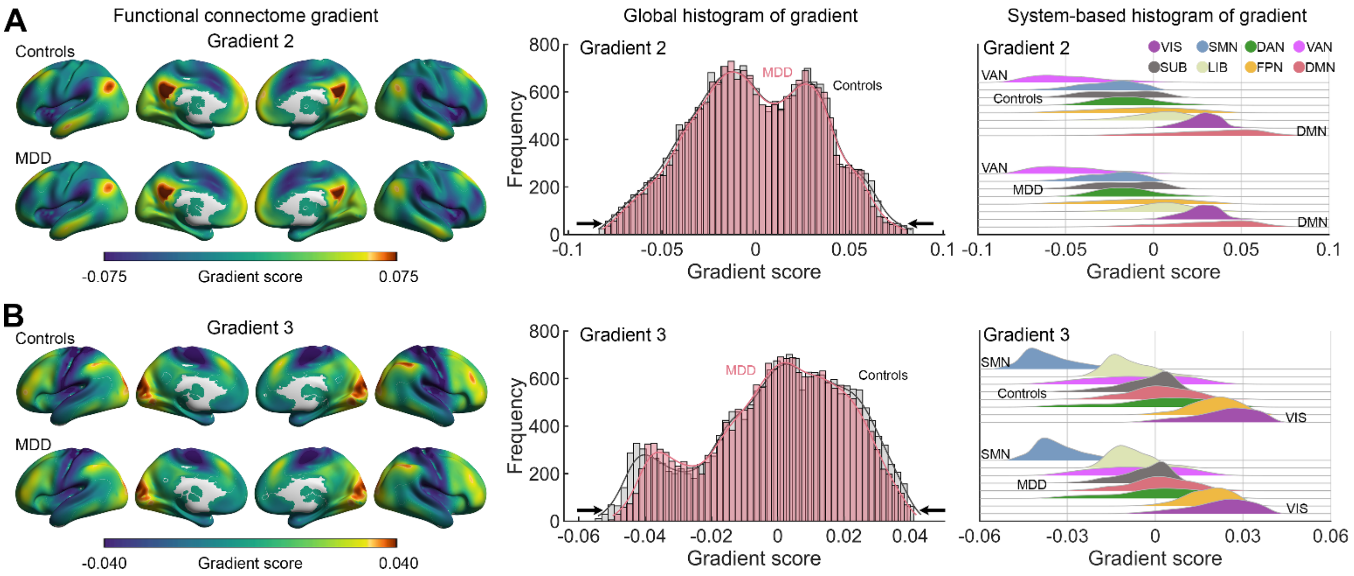


**Figure S3.** Connectome gradient mapping in patients with MDD and controls. **(A)** The second gradient extended between the default mode and the ventral attention networks. **(B)** The third gradient separated the sensorimotor from the visual networks. Global and system-based histograms show that the extreme values were contracted in patients with MDD relative to the controls for both gradients. Surface rendering was generated using BrainNet Viewer (www.nitrc.org/projects/bnv/) (19) with the inflated cortical 32K surface (13). VIS, visual network; SMN, sensorimotor network; DAN, dorsal attention network; VAN, ventral attention network; SUB, subcortical regions; LIB, limbic network; FPN, frontoparietal network; DMN, default mode network.

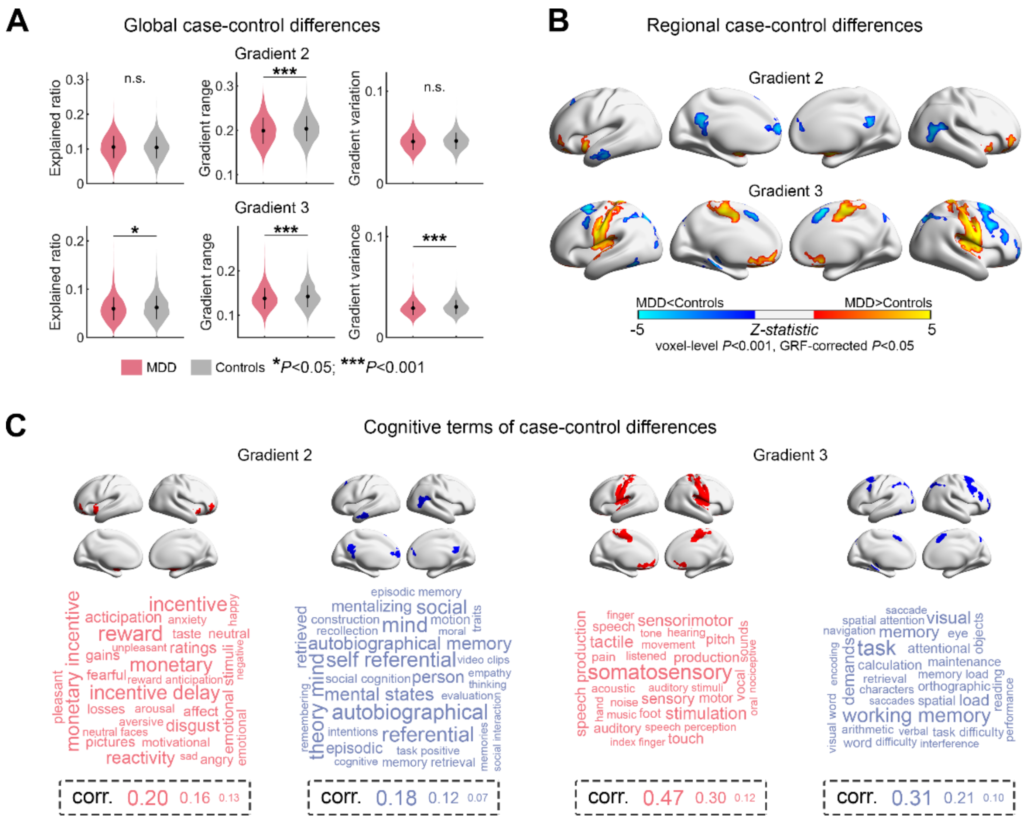


**Figure S4.** Statistical comparison of the second and third gradients between the MDD and control groups. **(A)** Case-control differences in global gradient metrics. *, $P < 0.05$; ***, $P < 0.001$; n.s., not significant. **(B)** Case-control differences in regional gradient scores, with higher/lower values in MDD presented as warm/cold colors. The significance level was set as voxel-level $P < 0.001$ and Gaussian random field cluster level-corrected $P < 0.05$. **(C)** Word clouds of cognitive functions associated with brain regions that exhibited higher (red) or lower (blue) gradient scores in MDD. The font size of the cognitive terms corresponds to the correlation of corresponding meta-analytic maps generated by Neurosynth.

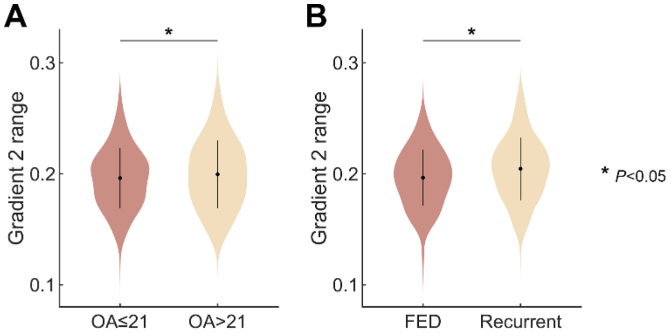


**Figure S5.** Effects of clinical factors. The range of the second gradient was significantly lower **(A)** in patients with an onset age ≤ 21 years than in those with an onset age older than 21 years and **(B)** in patients in their first episode than in recurrent patients, respectively. OA, onset age; FED, first episode depression.

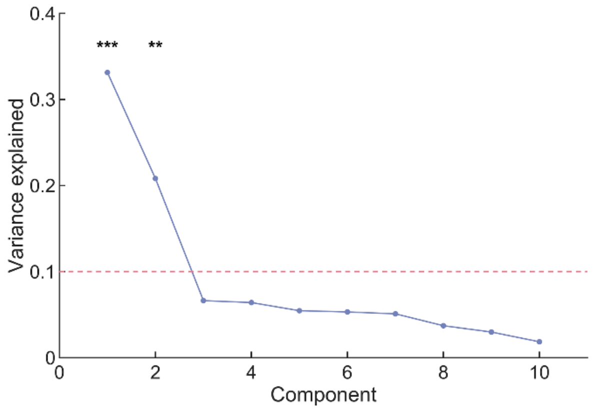


**Figure S6.** The percentage of variance in response variables explained by components in the partial least square regression analysis. The significance level was determined by a permutation test (N = 10,000) with spatial autocorrelation corrected. ***, $P < 0.001$; **, $P < 0.01$.

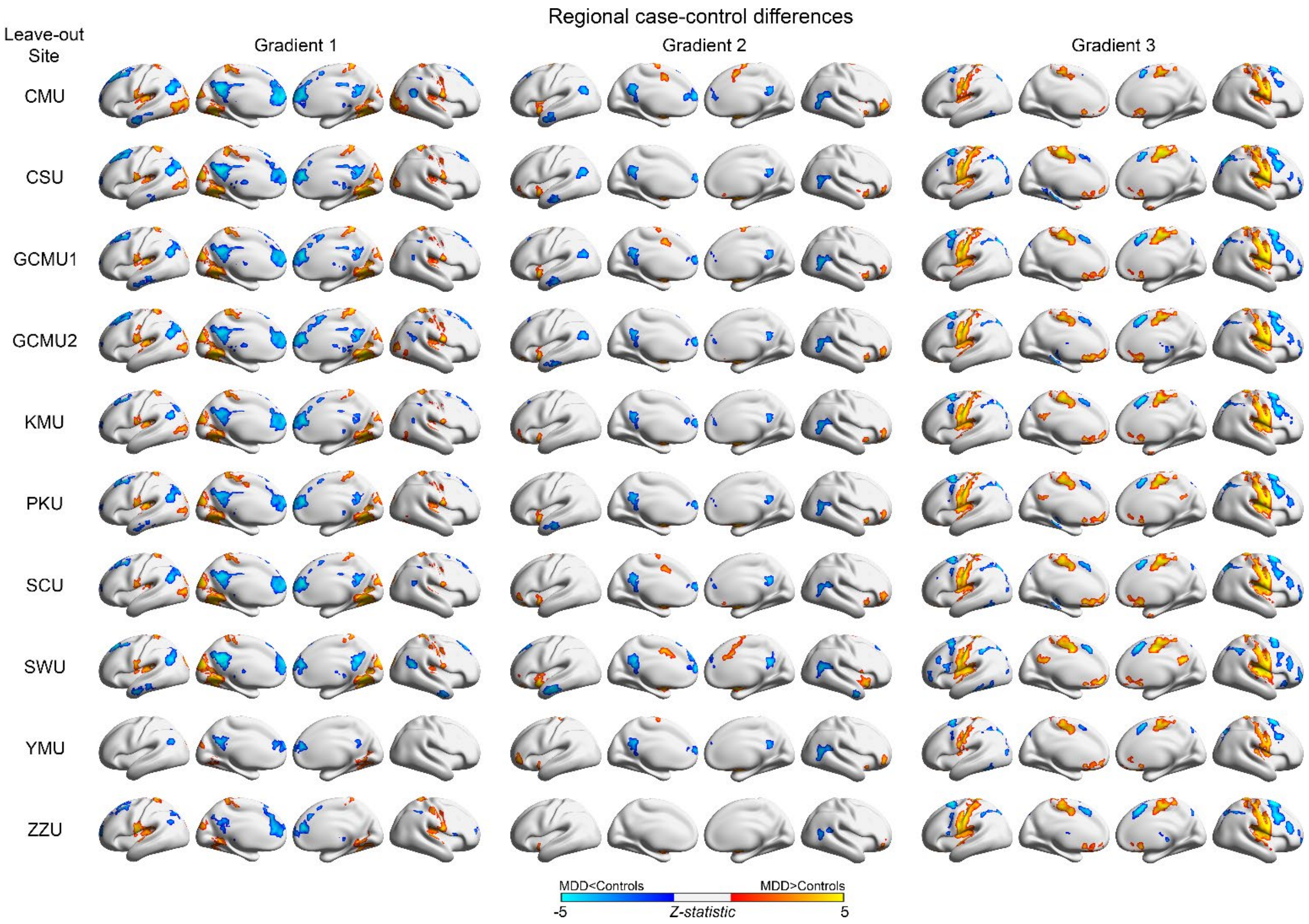


**Figure S7.** Leave-one-site-out cross-validation on regional alterations in the three gradient components. The case-control differences in regional gradient scores were largely comparable when excluding data from different centers and were highly similar to the main findings. Specifically, the MDD group showed significantly altered gradient scores compared with the control group (voxel-level $P < 0.001$, Gaussian random field cluster level-corrected $P < 0.05$): (*i*) in Gradient 1, lower gradient scores in the default mode network but higher scores in the visual and sensorimotor networks; (*ii*) in Gradient 2, lower scores in the default mode network but higher scores in the ventral attention network and subcortical regions; and (*iii*) in Gradient 3, lower scores in the frontoparietal network but higher scores in the sensorimotor network.

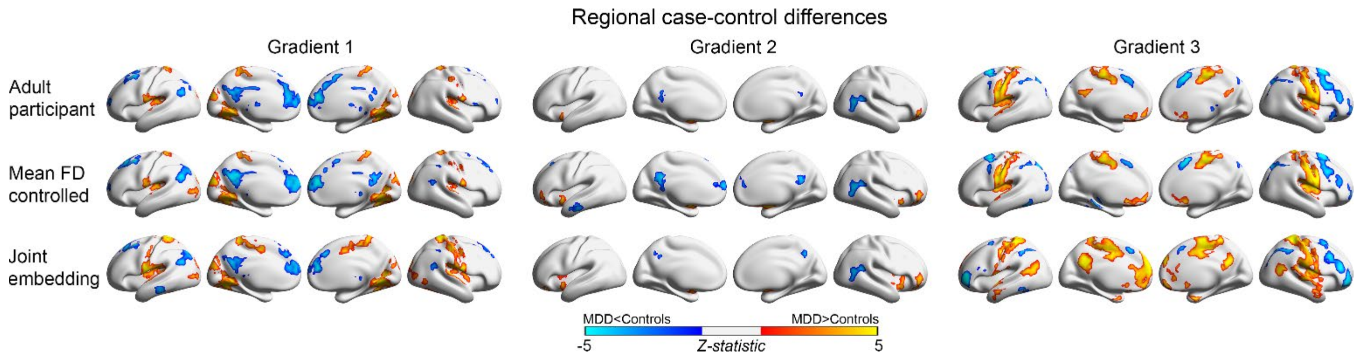


**Figure S8.** Gradient alterations derived from data from adult participants, statistical analysis controlling for mean framewise displacement (FD), and joint embedding alignment. The case-control differences in regional gradient scores were highly similar to the main findings in data from only adult participants when adding mean framewise displacement as an additional covariate in statistical analysis or using joint embedding in aligning individual gradient maps. Specifically, compared with the control group, the MDD group showed significantly altered gradient scores (voxel-level $P < 0.001$, Gaussian random field cluster level-corrected $P < 0.05$): (*i*) in Gradient 1, lower gradient scores in the default mode network but higher scores in the visual and sensorimotor networks; (*ii*) in Gradient 2, lower scores in the default mode network but higher scores in the ventral attention network and subcortical regions; and (*iii*) in Gradient 3, lower scores in the frontoparietal network but higher scores in the sensorimotor network. The default mode regions exhibited significantly higher scores of Gradient 3 in the patient group than in the control group, when joint embedding alignment was performed.

### Supplementary Tables

#### Table S1. Scan parameters of R-fMRI data in each center

| Center | Scanner | TR (ms) | TE (ms) | FA (°) | FOV ($mm^2$) | Matrix | Resolution ($mm^2$) | Slices | Thickness (mm) | Gap (mm) | VOL |
|---|---|---|---|---|---|---|---|---|---|---|---|
| CMU | GE HDxT 3T | 2000 | 40 | 90 | 240×240 | 64×64 | 3.75×3.75 | 35 | 3 | 0 | 200 |
| CSU | GE HDxT 3T | 2000 | 30 | 90 | 220×220 | 64×64 | 3.44×3.44 | 33 | 4 | 0.6 | 180 |
| GCMU1 | GE HDxT 3T | 2000 | 30 | 90 | 220×220 | 64×64 | 3.44×3.44 | 36 | 3 | 1 | 185 |
| GCMU2 | GE HDxT 3T | 2000 | 30 | 90 | 240×240 | 64×64 | 3.75×3.75 | 33 | 4 | 0 | 250 |
| KMU | PHILIPS Achieva 3T | 2200 | 35 | 90 | 230×230 | 128×128 | 1.80×1.80 | 50 | 3 | 0 | 240 |
| PKU | Siemens Trio 3T | 2000 | 30 | 90 | 210×210 | 64×64 | 3.28×3.28 | 30 | 4 | 0.8 | 210 |
| SCU | GE EXCITE 3T | 2000 | 30 | 90 | 220×220 | 64×64 | 3.44×3.44 | 30 | 5 | 0 | 200 |
| SWU | Siemens Trio 3T | 2000 | 30 | 90 | 220×220 | 64×64 | 3.44×3.44 | 32 | 3 | 1 | 242 |
| YMU | Siemens Trio 3T | 2500 | 27 | 77 | 220×220 | 64×64 | 3.44×3.44 | 43 | 3.4 | 0 | 200 |
| ZZU | GE MR750 3T | 2000 | 40 | 90 | 220×220 | 64×64 | 3.44×3.44 | 32 | 4 | 0.5 | 180 |

Abbreviations: TR, repetition time; TE, echo time; FA, flip angle; FOV, field of view; VOL, volume; CMU, China Medical University; CSU, Central South University; GCMU, Guangzhou University of Chinese Medicine; KMU, Kunming Medical University; PKU, Peking University; SCU, Sichuan University; SWU, Southwest University; YMU, National Yang-Ming University; ZZU, Zhengzhou University; GE, General Electric.

**Table S2. Case-control differences in global gradient metrics**

| Metric | MDD (N=1150), mean (SD) | Controls (N=1084), mean (SD) | *t* | Cohen's *d* | *P* |
|---|---|---|---|---|---|
| G1 explained ratio | 0.117 (0.031) | 0.121 (0.030) | -3.74 | -0.16 | 0.00019 |
| G2 explained ratio | 0.106 (0.032) | 0.104 (0.031) | 0.94 | 0.04 | 0.350 |
| G3 explained ratio | 0.060 (0.032) | 0.062 (0.024) | -2.34 | -0.10 | 0.019 |
| G1 range | 0.217 (0.036) | 0.224 (0.035) | -5.04 | -0.21 | 0.000001 |
| G2 range | 0.199 (0.029) | 0.204 (0.028) | -3.65 | -0.15 | 0.00027 |
| G3 range | 0.138 (0.024) | 0.142 (0.024) | -4.40 | -0.19 | 0.000011 |
| G1 variance | 0.050 (0.011) | 0.052 (0.011) | -4.81 | -0.20 | 0.000002 |
| G2 variance | 0.045 (0.009) | 0.046 (0.009) | -1.90 | -0.08 | 0.058 |
| G3 variance | 0.029 (0.007) | 0.030 (0.007) | -4.75 | -0.20 | 0.000002 |

Abbreviations: MDD, major depressive disorder; SD, standard deviation; G1, Gradient 1; G2, Gradient 2; G3, Gradient 3.

**Table S3. Clusters with significant between-group differences in the second gradient**

| No. | Region | x | y | z | *t* | Cohen's *d* | Size ($mm^3$) |
|---|---|---|---|---|---|---|---|
| MDD > Controls | | | | | | | |
| 1 | Right pallidum/putamen | 16 | 8 | 0 | 6.66 | 0.28 | 14336 |
| 2 | Left putamen | -28 | -8 | -4 | 6.27 | 0.27 | 12480 |
| 3 | Right inferior frontal gyrus, triangular/orbital parts, BA10 | 52 | 44 | 4 | 5.09 | 0.22 | 5760 |
| 4 | Left inferior frontal gyrus, orbital part, BA47 | -28 | 32 | -12 | 4.80 | 0.20 | 2752 |
| MDD < Controls | | | | | | | |
| 5 | Left posterior cingulate gyrus/precuneus, BA31 | -4 | -48 | 28 | -4.92 | -0.21 | 6400 |
| 6 | Left superior frontal gyrus, medial part, BA10 | -12 | 60 | 12 | -5.28 | -0.22 | 5184 |
| 7 | Right middle temporal gyrus, BA39 | 48 | -56 | 12 | -4.95 | -0.21 | 3840 |
| 8 | Left middle temporal gyrus, BA21 | -60 | -8 | -20 | -4.52 | -0.19 | 2432 |
| 9 | Left superior/middle frontal gyri, BA8 | -12 | 24 | 56 | -3.96 | -0.17 | 2368 |

**Table S4. Clusters with significant between-group differences in the third gradient**

| No. | Region | x | y | z | *t* | Cohen's *d* | Size ($mm^3$) |
|---|---|---|---|---|---|---|---|
| MDD > Controls | | | | | | | |
| 1 | Bilateral postcentral/precentral gyri, BA6/3/4 | -44 | -16 | 36 | 6.52 | 0.28 | 93504 |
| 2 | Left rectus gyrus, BA11 | -12 | 40 | -20 | 5.51 | 0.23 | 12288 |
| 3 | Right rectus gyrus, BA25 | 12 | 24 | -12 | 5.06 | 0.21 | 6720 |
| MDD < Controls | | | | | | | |
| 4 | Right supplementary motor area/middle frontal gyrus, BA6 | 8 | 12 | 56 | -6.10 | -0.26 | 31360 |
| 5 | Left inferior parietal gyrus, BA40 | -44 | -44 | 36 | -5.12 | -0.22 | 11200 |
| 6 | Right superior parietal gyri, BA7 | 32 | -72 | 48 | -5.04 | -0.21 | 9600 |
| 7 | Right middle frontal gyrus, orbital parts, BA10 | 32 | 52 | -4 | -4.65 | -0.20 | 2496 |
| 8 | Left middle frontal gyrus/inferior frontal gyrus, triangular part, BA9 | -52 | 20 | 40 | -4.33 | -0.18 | 1984 |
| 9 | Left inferior temporal/fusiform gyri, BA37 | -44 | -56 | -12 | -4.47 | -0.19 | 1728 |
| 10 | Left hippocampus | -16 | -28 | -12 | -4.84 | -0.20 | 1600 |

**Table S5. Enrichment of genes associated with MDD-related alterations in the principal gradient**

| GO Term | Description | *P* | FDR *q* | Enrichment | Genes |
|---|---|---|---|---|---|
| **Biological Process** | | | | | |
| **GO:0099536** | Synaptic signaling | $2.65\times10^{-7}$ | $3.63\times10^{-3}$ | 2.54 | CPNE6, TAC1, GLRA2, SNCA, SYT4, SST, GABRB1, NTSR1, HTR7, SYT10, HRH1, GRID2, HTR1A, EFNB3, CNR1, KCNMB4, CNIH2, GRIA1, PCDH8, RIT2, CBLN1, GABRA2, GRM1, GLRA3, GABRA5, SLC1A4, GABRA3, PRKCG, CNIH3, CBLN2, SLC17A7, DTNB |
| **GO:0099537** | Trans-synaptic signaling | $5.51\times10^{-7}$ | $3.78\times10^{-3}$ | 2.51 | CPNE6, TAC1, GLRA2, SNCA, SYT4, GABRB1, SST, NTSR1, HTR7, SYT10, HRH1, GRID2, HTR1A, EFNB3, CNR1, KCNMB4, CNIH2, GRIA1, PCDH8, RIT2, CBLN1, GABRA2, GRM1, GLRA3, GABRA5, SLC1A4, GABRA3, PRKCG, CNIH3, CBLN2, SLC17A7 |
| **GO:0098916** | Anterograde trans-synaptic signaling | $1.02\times10^{-6}$ | $4.65\times10^{-3}$ | 2.52 | CPNE6, TAC1, GLRA2, SNCA, GABRB1, SST, HTR7, NTSR1, SYT10, HRH1, GRID2, HTR1A, KCNMB4, CNIH2, GRIA1, PCDH8, RIT2, CBLN1, GABRA2, GRM1, GLRA3, GABRA5, GABRA3, SLC1A4, PRKCG, CNIH3, CBLN2, SLC17A7 |
| **GO:0007268** | Chemical synaptic transmission | $1.02\times10^{-6}$ | $3.48\times10^{-3}$ | 2.52 | CPNE6, TAC1, GLRA2, SNCA, SST, GABRB1, HTR7, NTSR1, SYT10, HRH1, GRID2, HTR1A, KCNMB4, CNIH2, GRIA1, PCDH8, RIT2, CBLN1, GABRA2, GRM1, GLRA3, GABRA5, GABRA3, SLC1A4, PRKCG, CBLN2, SLC17A7 |
| **GO:0007267** | Cell-cell signaling | $9.61\times10^{-6}$ | $2.63\times10^{-2}$ | 2.06 | CPNE6, TAC1, GLRA2, SNCA, SYT4, NRP1, SST, GABRB1, NTSR1, HTR7, SYT10, HRH1, GRID2, EFNB3, HTR1A, CNR1, KCNMB4, FZD1, CNIH2, GRIA1, PCDH8, GPNMB, RIT2, CBLN1, GABRA2, GRM1, GLRA3, GABRA5, SLC1A4, GABRA3, SEMA4F, PRKCG, CNIH3, CBLN2, ADRA1B, SLC17A7, DTNB |
| **Molecular Function** | | | | | |
| **GO:0005509** | Calcium ion binding | $7.49\times10^{-6}$ | $2.92\times10^{-2}$ | 1.90 | MAN1A1, CAPNS1, PCDH15, SCUBE2, F12, NKD2, CAPN12, SYT5, SYT17, SYT4, SCGN, EFEMP2, CASQ1, MAN1B1, SYT10, EFCAB1, PCDH8, CALB1, PCDH10, VWCE, FSTL5, AIF1, HPCAL1, PAMR1, PLCB2, MYL5, CALM3, CCBE1, IDS, LTBP4, CPNE6, HPCA, PCDH20, PPEF1, MYL12B, PLCH1, SNCA, CETN2, PLCH2, CABYR, HPCAL4, ACTN2, DLL3, CGREF1, GNPTAB, CIB2, CIB1, NECAB2, SUSD1, PCDH17, SULF1, S100A6, EFHC2, SLIT1, C1S, SULF2, RASGRP1, DOC2A, CDH4, DOC2B, CDH10, CD248, CDH11, CDH8, CDH9, CDH13, NCAN, FBLN2, PCDH19, BAIAP3 |

**Table S6. Differences in global gradient metrics between MDD subgroups**

| Metric | First-episode vs. Recurrent | Nonmedicated vs. Medicated | Onset Age>21 vs. Onset Age≤21 |
|---|---|---|---|
| Sample Size (N) | First-episode=512<br>Recurrent=80 | Nonmedicated=622<br>Medicated=277 | Onset Age>21=293<br>Onset Age≤21=303 |
| G1 explained ratio | *t* =0.30; *P*=0.77; *d*=0.04 | *t*=-1.90; *P*=0.058; *d*=-0.14 | *t*=1.69; *P*=0.091; *d*=0.14 |
| G2 explained ratio | *t*=-0.95; *P*=0.34; *d*=-0.11 | *t*=1.87; *P*=0.062; *d*=0.14 | *t*=1.16; *P*=0.25; *d*=0.14 |
| G3 explained ratio | *t*=0.80; *P*=0.94; *d*=0.10 | *t*=-0.45; *P*=0.65; *d*=-0.03 | *t*=-0.53; *P*=0.60; *d*=-0.04 |
| G1 range | *t*=-0.87; *P*=0.39; *d*=-0.10 | *t*=-0.63; *P*=0.53; *d*=-0.05 | ***t*=3.38; *P*=0.001; *d*=0.28** |
| G2 range | ***t*=-2.22; *P*=0.027; *d*=-0.27** | *t*=1.24; *P*=0.21; *d*=0.09 | ***t*=2.54; *P*=0.011; *d*=0.21** |
| G3 range | *t*=-0.47; *P*=0.64; *d*=-0.06 | *t*=-0.63; *P*=0.53; *d*=-0.05 | *t*=0.75; *P*=0.45; *d*=0.06 |
| G1 variance | *t*=-0.62; *P*=0.54; *d*=-0.07 | *t*=-1.55; *P*=0.12; *d*=-0.11 | ***t*=2.23; *P*=0.026; *d*=0.18** |
| G2 variance | *t*=-1.51; *P*=0.13; *d*=-0.18 | *t*=1.32; *P*=0.19; *d*=0.10 | *t*=-1.96; *P*=0.050; *d*=-0.16 |
| G3 variance | *t*=1.14; *P*=0.25; *d*=0.14 | *t*=-0.50; *P*=0.62; *d*=-0.04 | *t*=-0.55; *P*=0.58; *d*=0.05 |

G1, Gradient 1; G2, Gradient 2; G3, Gradient 3.

**Table S7. Leave-one-site-out cross-validation on alterations in global gradient metrics**

| Leave-out site | Sample Size (N) | G1 explained ratio | G2 explained ratio | G3 explained ratio | G1 range | G2 range | G3 range | G1 variance | G2 variance | G3 variance |
|---|---|---|---|---|---|---|---|---|---|---|
| CMU | MDD=1025 | ***t*=-4.05** | *t*=-0.46 | *t*=-1.82 | ***t*=-5.29** | ***t*=-3.72** | ***t*=-3.35** | ***t*=-5.11** | ***t*=-2.18** | ***t*=-3.18** |
| | Controls=835 | ***P*<0.001** | *P*=0.648 | *P*=0.069 | ***P*<0.001** | ***P*<0.001** | ***P*=0.001** | ***P*<0.001** | ***P*=0.029** | ***P*=0.001** |
| | | ***d*=-0.19** | *d*=-0.02 | *d*=-0.08 | ***d*=-0.25** | ***d*=-0.17** | ***d*=-0.16** | ***d*=-0.24** | ***d*=-0.10** | ***d*=-0.15** |
| CSU | MDD=973 | ***t*=-4.06** | *t*=-0.93 | *t*=-1.97 | ***t*=-5.33** | ***t*=-3.24** | ***t*=-4.62** | ***t*=-5.17** | *t*=-1.80 | ***t*=-4.94** |
| | Controls=976 | ***P*<0.001** | *P*=0.351 | *P*=0.050 | ***P*<0.001** | ***P*=0.001** | ***P*<0.001** | ***P*<0.001** | *P*=0.072 | ***P*<0.001** |
| | | ***d*=-0.18** | *d*=-0.04 | *d*=-0.09 | ***d*=-0.24** | ***d*=-0.15** | ***d*=-0.21** | ***d*=-0.23** | *d*=-0.08 | ***d*=-0.22** |
| GCMU_1 | MDD=1116 | ***t*=-3.68** | *t*=0.89 | ***t*=-2.15** | ***t*=-4.88** | ***t*=-3.67** | ***t*=-4.41** | ***t*=-4.63** | *t*=-1.88 | ***t*=-4.69** |
| | Controls=1050 | ***P*<0.001** | *P*=0.372 | ***P*=0.032** | ***P*<0.001** | ***P*=0.001** | ***P*<0.001** | ***P*<0.001** | *P*=0.061 | ***P*<0.001** |
| | | ***d*=-0.16** | *d*=0.04 | ***d*=-0.09** | ***d*=-0.21** | ***d*=-0.16** | ***d*=-0.19** | ***d*=-0.20** | *d*=-0.08 | ***d*=-0.20** |
| GCMU_2 | MDD=1084 | ***t*=-4.05** | *t*=1.10 | ***t*=-2.30** | ***t*=-5.17** | ***t*=-3.47** | ***t*=-4.11** | ***t*=-4.97** | *t*=-1.72 | ***t*=-4.55** |
| | Controls=1018 | ***P*<0.001** | *P*=0.27 | ***P*=0.021** | ***P*<0.001** | ***P*=0.001** | ***P*<0.001** | ***P*<0.001** | *P*=0.086 | ***P*<0.001** |
| | | ***d*=-0.18** | *d*=0.05 | ***d*=-0.10** | ***d*=-0.23** | ***d*=-0.15** | ***d*=-0.18** | ***d*=-0.22** | *d*=-0.08 | ***d*=-0.20** |
| KMU | MDD=1109 | ***t*=-3.87** | *t*=0.86 | ***t*=-1.99** | ***t*=-4.86** | ***t*=-3.51** | ***t*=-4.26** | ***t*=-4.67** | *t*=-1.90 | ***t*=-4.56** |
| | Controls=1034 | ***P*<0.001** | *P*=0.391 | ***P*=0.047** | ***P*<0.001** | ***P*<0.001** | ***P*<0.001** | ***P*<0.001** | *P*=0.058 | ***P*<0.001** |
| | | ***d*=-0.17** | *d*=0.04 | ***d*=-0.09** | ***d*=-0.21** | ***d*=-0.15** | ***d*=-0.18** | ***d*=-0.20** | *d*=-0.08 | ***d*=-0.20** |
| PKU | MDD=1075 | ***t*=-3.77** | *t*=-0.95 | ***t*=-2.14** | ***t*=-4.70** | ***t*=-3.82** | ***t*=-4.06** | ***t*=-4.70** | *t*=-1.96 | ***t*=-4.50** |
| | Controls=1011 | ***P*<0.001** | *P*=0.344 | ***P*=0.033** | ***P*<0.001** | ***P*<0.001** | ***P*<0.001** | ***P*<0.001** | *P*=0.050 | ***P*<0.001** |
| | | ***d*=-0.17** | *d*=-0.04 | ***d*=-0.09** | ***d*=-0.21** | ***d*=-0.17** | ***d*=-0.18** | ***d*=-0.21** | *d*=-0.09 | ***d*=-0.20** |
| SCU | MDD=1100 | ***t*=-3.43** | *t*=-0.77 | ***t*=-2.24** | ***t*=-4.85** | ***t*=-3.46** | ***t*=-4.30** | ***t*=-4.47** | *t*=-1.75 | ***t*=-4.62** |
| | Controls=1043 | ***P*=0.001** | *P*=0.442 | ***P*=0.025** | ***P*<0.001** | ***P*=0.001** | ***P*<0.001** | ***P*<0.001** | *P*=0.080 | ***P*<0.001** |
| | | ***d*=-0.15** | *d*=-0.03 | ***d*=-0.10** | ***d*=-0.21** | ***d*=-0.15** | ***d*=-0.19** | ***d*=-0.19** | *d*=-0.08 | ***d*=-0.17** |
| SWU | MDD=868 | ***t*=-2.94** | *t*=-0.02 | ***t*=-3.27** | ***t*=-5.23** | ***t*=-4.32** | ***t*=-4.74** | ***t*=-4.78** | ***t*=-2.91** | ***t*=-5.33** |
| | Controls=830 | ***P*=0.003** | *P*=0.987 | ***P*=0.001** | ***P*<0.001** | ***P*<0.001** | ***P*<0.001** | ***P*<0.001** | ***P*=0.004** | ***P*<0.001** |
| | | ***d*=-0.14** | *d*=0.00 | ***d*=-0.16** | ***d*=-0.25** | ***d*=-0.21** | ***d*=-0.23** | ***d*=-0.23** | ***d*=-0.14** | ***d*=-0.26** |
| YMU | MDD=1045 | ***t*=-2.23** | *t*=-0.72 | *t*=-1.97 | ***t*=-3.99** | ***t*=-3.27** | ***t*=-3.68** | ***t*=-3.12** | *t*=-1.62 | ***t*=-3.83** |
| | Controls=975 | ***P*=0.033** | *P*=0.474 | *P*=0.053 | ***P*<0.001** | ***P*=0.001** | ***P*<0.001** | ***P*=0.002** | *P*=0.105 | ***P*<0.001** |
| | | ***d*=-0.10** | *d*=-0.03 | *d*=-0.09 | ***d*=-0.18** | ***d*=-0.15** | ***d*=-0.16** | ***d*=-0.14** | *d*=-0.07 | ***d*=-0.17** |
| ZZU | MDD=955 | ***t*=-3.41** | ***t*=2.13** | *t*=-1.81 | ***t*=-3.34** | *t*=-1.81 | ***t*=-4.34** | ***t*=-3.94** | *t*=-0.26 | ***t*=-4.93** |
| | Controls=984 | ***P*=0.001** | ***P*=0.033** | *P*=0.070 | ***P*=0.001** | *P*=0.070 | ***P*<0.001** | ***P*<0.001** | *P*=0.792 | ***P*<0.001** |
| | | ***d*=-0.15** | ***d*=-0.10** | *d*=-0.08 | ***d*=-0.15** | *d*=-0.08 | ***d*=-0.20** | ***d*=-0.18** | *d*=-0.01 | ***d*=-0.22** |

G1, Gradient 1; G2, Gradient 2; G3, Gradient 3; CMU, China Medical University; CSU, Central South University; GCMU, Guangzhou University of Chinese Medicine; KMU, Kunming Medical University; PKU, Peking University; SCU, Sichuan University; SWU, Southwest University; YMU, National Yang-Ming University; ZZU, Zhengzhou University.

**Table S8. Reproducibility validation in adults and for head-motion effect**

| Metric | Adult participant | Control for mean FD | Joint embedding alignment |
|---|---|---|---|
| Sample Size (N) | MDD=1002; Controls=1034 | MDD=1150; Controls=1084 | MDD=1150; Controls=1084 |
| G1 explained ratio | ***t*=-3.40; *P*=0.001; *d*=-0.15** | ***t*=-3.77; *P*<0.001; *d*=-0.16** | ***t*=-3.38; *P*=0.001; *d*=-0.14** |
| G2 explained ratio | *t*=1.68; *P*=0.093; *d*=0.07 | *t*=1.02; *P*=0.307; *d*=0.04 | *t*=-0.74; *P*=0.460; *d*=-0.03 |
| G3 explained ratio | ***t*=-2.26; *P*=0.024; *d*=-0.10** | ***t*=-2.35; *P*=0.019; *d*=-0.10** | *t*=-0.63; *P*=0.531; *d*=-0.03 |
| G1 range | ***t*=-3.99; *P*<0.001; *d*=-0.18** | ***t*=-5.01; *P*<0.001; *d*=-0.21** | ***t*=-4.63; *P*<0.001; *d*=-0.20** |
| G2 range | ***t*=-2.24; *P*=0.026; *d*=-0.10** | ***t*=-3.56; *P*<0.001; *d*=-0.15** | ***t*=-7.18; *P*<0.001; *d*=-0.30** |
| G3 range | ***t*=-4.11; *P*<0.001; *d*=-0.18** | ***t*=-4.48; *P*<0.001; *d*=-0.19** | ***t*=-3.46; *P*=0.001; *d*=-0.15** |
| G1 variance | ***t*=-4.26; *P*<0.001; *d*=-0.19** | ***t*=-4.76; *P*<0.001; *d*=-0.20** | ***t*=-5.86; *P*<0.001; *d*=-0.25** |
| G2 variance | *t*=-0.56; *P*=0.574; *d*=-0.02 | *t*=-1.81; *P*=0.071; *d*=-0.08 | ***t*=-6.58; *P*<0.001; *d*=-0.28** |
| G3 variance | ***t*=-4.59; *P*<0.001; *d*=-0.20** | ***t*=-4.81; *P*<0.001; *d*=-0.20** | ***t*=-3.16; *P*=0.002; *d*=-0.13** |

G1, Gradient 1; G2, Gradient 2; G3, Gradient 3; FD, framewise displacement.
